# Enterobacterial repetitive intergenic consensus (ERIC)-PCR analysis as a trace for *Burkholderia pseudomallei* in Myanmar

**DOI:** 10.1101/2024.02.15.580599

**Authors:** Nay Myo Aung, Khine Khine Su, Narisara Chantratita, Chanwit Tribuddharat

**Author notes:** **Corresponding author:** Mailing address: Department of Microbiology, Defense Services Medical Academy, 11021, Myanmar.

## Abstract

Melioidosis is a potentially fatal disease caused by *Burkholderia pseudomallei*, which is endemic in Southeast Asia, including Myanmar. The typeability of enterobacterial repetitive intergenic consensus (ERIC)-PCR assessed for 21 *B. pseudomallei*, they used the results of sequence types (STs) of the multilocus sequence typing (MLST) method. Among 5 soil and 16 clinical *B. pseudomallei* isolates, the most significant bands were similar in position but different in minor band formation. ST 90 of two soil strains (Tontae_NMBP001 and Tontae_NMBP002) displayed the same ERIC banding pattern, while ST 56 of two clinical isolates (MMBP005 and MMBP010) from different regions exhibited a single type. The same ST found both clusters in the MLST method. The shared group STs showed four or three satellite variants in the MLST scheme. One novel studied ST (ST 1729) and regarded it as an out-group in the ERIC pattern. ERIC PCR demonstrated high discriminatory power, while MLST provided more discrimination for genetic diversity. MLST requires extensive sequencing and bioinformatics analysis, making it challenging to implement in resource-limited settings. More isolates are needed to validate these findings. Despite its limitations, ERIC PCR represents a valuable and cost-effective alternative to MLST for molecular typing of B. pseudomallei in resource-limited settings.

## Introduction

Melioidosis is an infectious disease caused by the gram-negative bacterium *Burkholderia pseudomallei*, which is prevalent in the soil and water of Southeast Asia and Northern Australia. The bacterium is an opportunistic pathogen that can cause a wide range of clinical manifestations, from acute sepsis to chronic infections, with mortality rates as high as 40% (1). Early diagnosis and prompt treatment with appropriate antibiotics are crucial for successful outcomes; however, the accurate identification and typing of *B. pseudomallei* remains challenging, particularly in resource-limited settings.

Molecular techniques have emerged as valuable tools for identifying and typing *B. pseudomallei* isolates. Among these, enterobacterial repetitive intergenic consensus polymerase chain reaction (ERIC PCR) and multilocus sequence typing (MLST) has widely used for the molecular epidemiology and phylogenetic analysis of *B. pseudomallei* (2). ERIC PCR is a PCR-based technique that amplifies the repetitive elements within the bacterial genome, producing a DNA fingerprint that can use for strain typing and clustering analysis (3). MLST, on the other hand, is a sequence-based method that targets specific genes in the bacterial genome, enabling the identification of unique alleles and the determination of genetic relatedness among isolates (4).

Despite the usefulness of MLST in identifying genetic variations and tracing the transmission of *B. pseudomallei*, its implementation can be problematic in resource-limited settings due to its high cost and technical requirements. In contrast, ERIC PCR is a simple and cost-effective alternative method for the molecular typing of bacteria, including *B. pseudomallei*. This technique amplifies the regions flanking the Enterobacterial Repetitive Intergenic Consensus (ERIC) sequence, a repetitive DNA element in multiple copies in bacterial genomes. The resulting banding patterns can be analyzed using gel electrophoresis, and clusters of strains with similar patterns can be identified.

However, while ERIC PCR is a helpful tool for molecular epidemiology studies, it has some limitations. For instance, it may not be as reliable as MLST in identifying genetically closely related strains, as it depends on intergenic regions’ variability rather than specific nucleotide changes. Additionally, interpreting ERIC PCR results can be subjective, as the banding patterns can be affected by experimental conditions and the interpretation of gel images (5). Despite its limitations, ERIC PCR represents a valuable and cost-effective alternative to MLST for molecular typing of *B. pseudomallei* in resource-limited settings. Its simplicity and low cost could be available for surveillance and outbreak investigations, particularly in endemic areas with limited advanced molecular methods.

In the context of Myanmar, where melioidosis is endemic, using these molecular techniques to identify and type B. pseudomallei is essential for epidemiological investigations and surveillance. However, the applicability of these methods in resource-limited settings needs to evaluate. This study aims to provide an overview of the use of ERIC PCR and MLST for identifying and typing B. pseudomallei in Myanmar and their potential as tools for the surveillance and control of melioidosis.

## Materials and Methods

### Bacterial strain collection

Five soil and sixteen clinical isolates of *Burkholderia pseudomallei* were collected in a previous study (6). Briefly, the published primers in the pudmlst website were used to amplify the published housekeeping gene fragments (*ace, gltB, gmhD, lepA, lipA, narK, ndh*) (19, 158). (https://pubmlst.org/bpseudomallei/). The PCR condition was evaluated in a previous study, and continued amplicon sequencing was done using Sanger methods (First Base company, Malaysia). Each isolate was analyzed by a string of seven integers (the allelic profile), which correspond to the allele numbers at the seven loci, in the order *ace-gltB-gmhD-lepA-lipA-narK-ndh*. Next, each unique allelic profile was considered a clone and was assigned a sequence type (ST), which also gave a convenient descriptor for the clone. An MLST database containing the sequences of all alleles, the allelic profiles, and information about the *B. pseudomallei* isolates, together with analysis tools, was recorded at Imperial College (London, United Kingdom) and can be examined on the *B. pseudomallei* pages of the MLST website (www.mlst.net). The resulting sequences at the seven loci were concatenated in the order of loci used to determine the allelic profile.

For the genotyping of *B. pseudomallei*, we performed ERIC-PCR again, and a pair of forward and reverse primers were used according to the reference article (3). The primers of 5’-ATG TAA GCT CCT GGG GAT TCA C-3’ (F) and 5’-AAG TAA GTG ACT GGG GTG AGC G-3’ (R) were applied. The reaction was performed in a volume of in 20 μl volumes containing 0.2 μl of 1 U of DNA polymerase (Thermoscience), 1 μl of DNA solution, 2 μl of 1x standard Taq reaction buffer (with MgCl2), 0.5 μl of 0.25 mM each dATP, dCTP, dGTP, and dTTP, and 0.5 μl of 0.5 μM each primer, adding 15.3 DNase free water. Finally, the thermocycler was programmed. Simultaneously, negative (*Burkholderia* species) were used to observe the results accurately. Gel bands of each isolate were examined under the installed software of the gel documentation system.

### Ethics review

The Siriraj Institutional Review Board approved the study (SIRB number: 546/2562 (EC1).

## Results

Among 21 isolates, ST 90 (n=6, 28.57%) was found as common ST from 3 clinical and soil isolates, respectively (Table 1). The remaining isolates were resulted as previously published and uploaded sequence types ST300 (n=1, 4.76%), ST 56 (n=2, 9.52%), ST 354 (n=2, 9.52%), ST 416 (n=1, 4.76%), which were isolated from clinical samples, whereas soil isolate showed ST 42 (n=1, 4.76%). The rest 8 isolates were identified in novel STs, representing ST 1722, ST 1723, ST 1724, ST 1725, ST 1727, ST 1728, and ST 1729 from clinical samples and ST 1726 from soil samples (Table 4.13).

**Table 1.** Myanmar *B. pseudomallei* isolates analyzed by multilocus sequence typing.

| Strain | Source | Type of specimen | Year | ST | Allele profile |  |  |  |  |  |  |
| --- | --- | --- | --- | --- | --- | --- | --- | --- | --- | --- | --- |
|  |  |  |  |  | *ace* | *gltB* | *gmhD* | *lepA* | *lipA* | *narK* | *ndh* |
| MMBP001 | Human | blood | 2018 | 300 | 1 | 1 | 3 | 1 | 1 | 4 | 1 |
| MMBP002 | Human | blood | 2018 | 1722<sup>a</sup> | 4 | 2 | 3 | 1 | 1 | 2 | 3 |
| MMBP003 | Human | Urine | 2018 | 1723<sup>a</sup> | 1 | 4 | 49 | 1 | 1 | 2 | 1 |
| MMBP004 | Human | wound | 2018 | 1728<sup>a</sup> | 1 | 12 | 6 | 1 | 10 | 4 | 1 |
| MMBP005 | Human | blood | 2018 | 56 | 3 | 1 | 4 | 1 | 1 | 4 | 1 |
| MMBP006 | Human | tissue | 2018 | 1724<sup>a</sup> | 1 | 1 | 3 | 1 | 8 | 2 | 1 |
| MMBP007 | Human | blood | 2018 | 1725<sup>a</sup> | 1 | 2 | 3 | 2 | 3 | 3 | 3 |
| MMBP008 | Human | blood | 2018 | 354 | 1 | 1 | 3 | 2 | 1 | 4 | 1 |
| MMBP009 | Human | blood | 2018 | 354 | 1 | 1 | 3 | 2 | 1 | 4 | 1 |
| MMBP010 | Human | blood | 2018 | 56 | 3 | 1 | 4 | 1 | 1 | 4 | 1 |
| MMBP011 | Human | Urine | 2018 | 1729<sup>a</sup> | 1 | 12 | 6 | 1 | 9 | 4 | 1 |
| MMBP012 | Human | blood | 2018 | 1727<sup>a</sup> | 1 | 12 | 13 | 2 | 1 | 1 | 3 |
| MMBP013 | Human | pleural fluid | 2018 | 416 | 1 | 12 | 6 | 1 | 1 | 4 | 1 |
| MMBP014 | Human | blood | 2018 | 90 | 1 | 12 | 6 | 1 | 1 | 4 | 1 |
| MMBP015 | Human | pus | 2018 | 90 | 1 | 12 | 6 | 1 | 1 | 4 | 1 |
| MMBP016 | Human | blood | 2018 | 90 | 1 | 1 | 6 | 2 | 1 | 42 | 1 |
| Tontae_NMBP001 | Soil | Soil | 2018 | 90 | 1 | 12 | 6 | 1 | 1 | 4 | 1 |
| Tontae_NMBP002 | Soil | Soil | 2018 | 90 | 1 | 12 | 6 | 1 | 1 | 4 | 1 |
| Tontae_NMBP003 | Soil | Soil | 2018 | 90 | 1 | 12 | 6 | 1 | 1 | 4 | 1 |
| Pathein_NMBP004 | Soil | Soil | 2018 | 42 | 1 | 12 | 6 | 2 | 1 | 2 | 1 |
| Pathein_NMBP005 | Soil | Soil | 2018 | 1726<sup>a</sup> | 1 | 10 | 6 | 2 | 1 | 2 | 1 |

<sup>a</sup> Showing novel ST

As a resource-limited country, Myanmar, the rapid, cost-effective, and flexible genotyping method for *B. pseudomallei* isolates was developed, presenting the Enterobacterial Repetitive Intergenic Consensus Polymerase Chain Reaction (ERIC-PCR) technique. Among 5 soil and 16 clinical *B. pseudomallei* isolates, it was seen that most of the major bands were quite similar in position but different in minor band formation. Therefore, ST 90 of two soil strains (Tontae_NMBP001 and Tontae_NMBP002) displayed the same ERIC banding pattern, while ST 56 of two clinical isolates (MMBP005 and MMBP010) exhibited a single type. Surprisingly, both of those two clusters were found to be the same ST in the MLST method (Figure. 1). It is noteworthy to reveal that both clinical isolates with ST 56 were obtained from patients residing in the same region of Yangon, which encompasses different cities such as Hlegu and Khayan (Figure. 2). Overall, ST 90 were approximately analyzed as a same clade, including one novel ST 1726. One novel ST (ST 1724) in this study was found in the same cluster with old published ST 300 in global data, showing DLV difference in the MLST scheme. It was observed that above mentioned 2 isolates exhibited 80% similarity in the ERIC pattern. However, another novel STs in this study shared the same groups with published STs (e.g., ST 1722 and ST 90, and ST 354 and ST 1725). The shared group STs showed four or three satellite variants in the MLST scheme. One novel studied ST (ST 1729) and was regarded as an out-group in the ERIC pattern.

**Figure 1.**
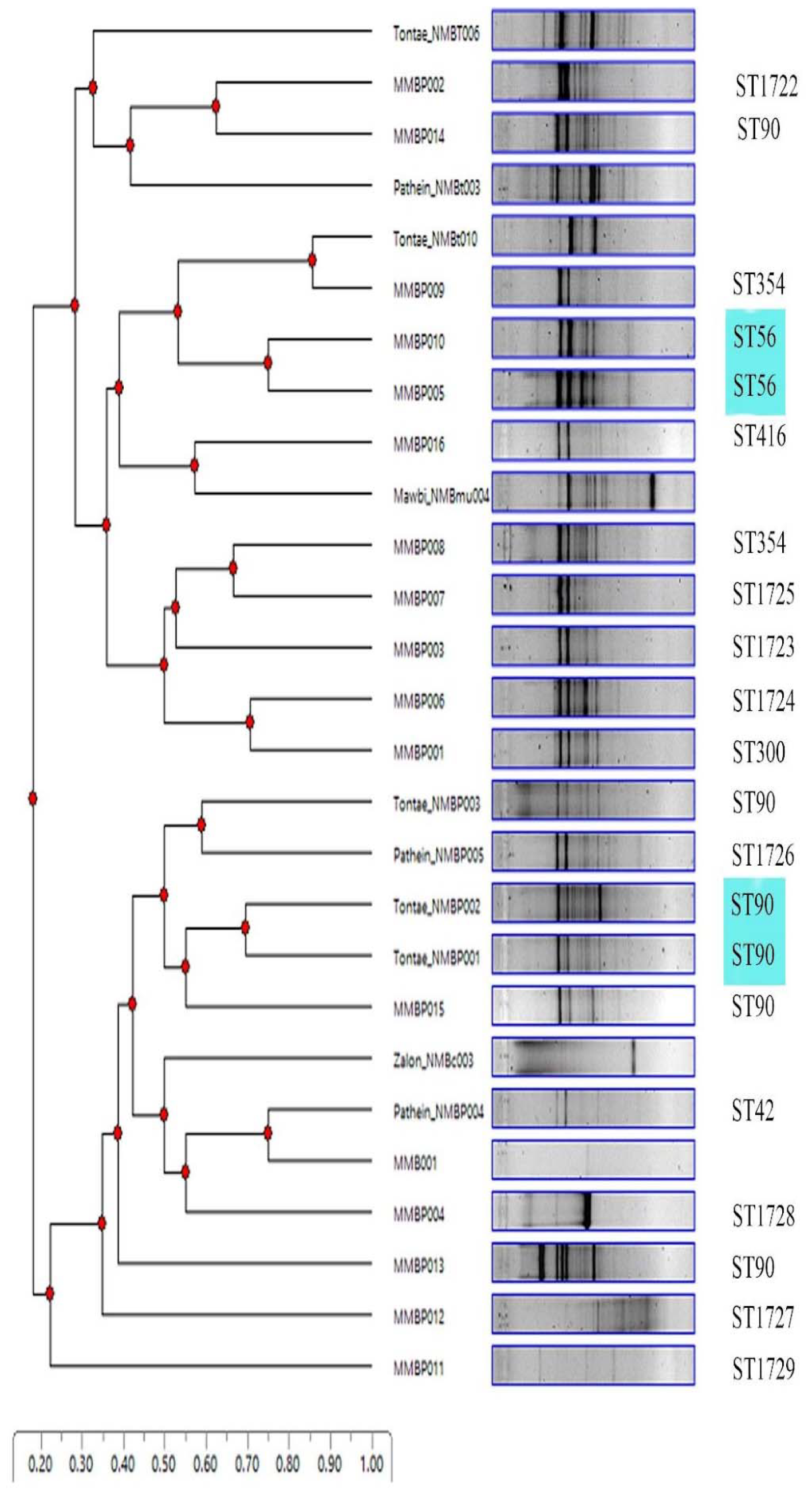
Assessment of ERIC patterns with related STs in Myanmar and highlight boxes showed the same patterns with the same STs. The same clone was a color-coded group

**Figure 2.**
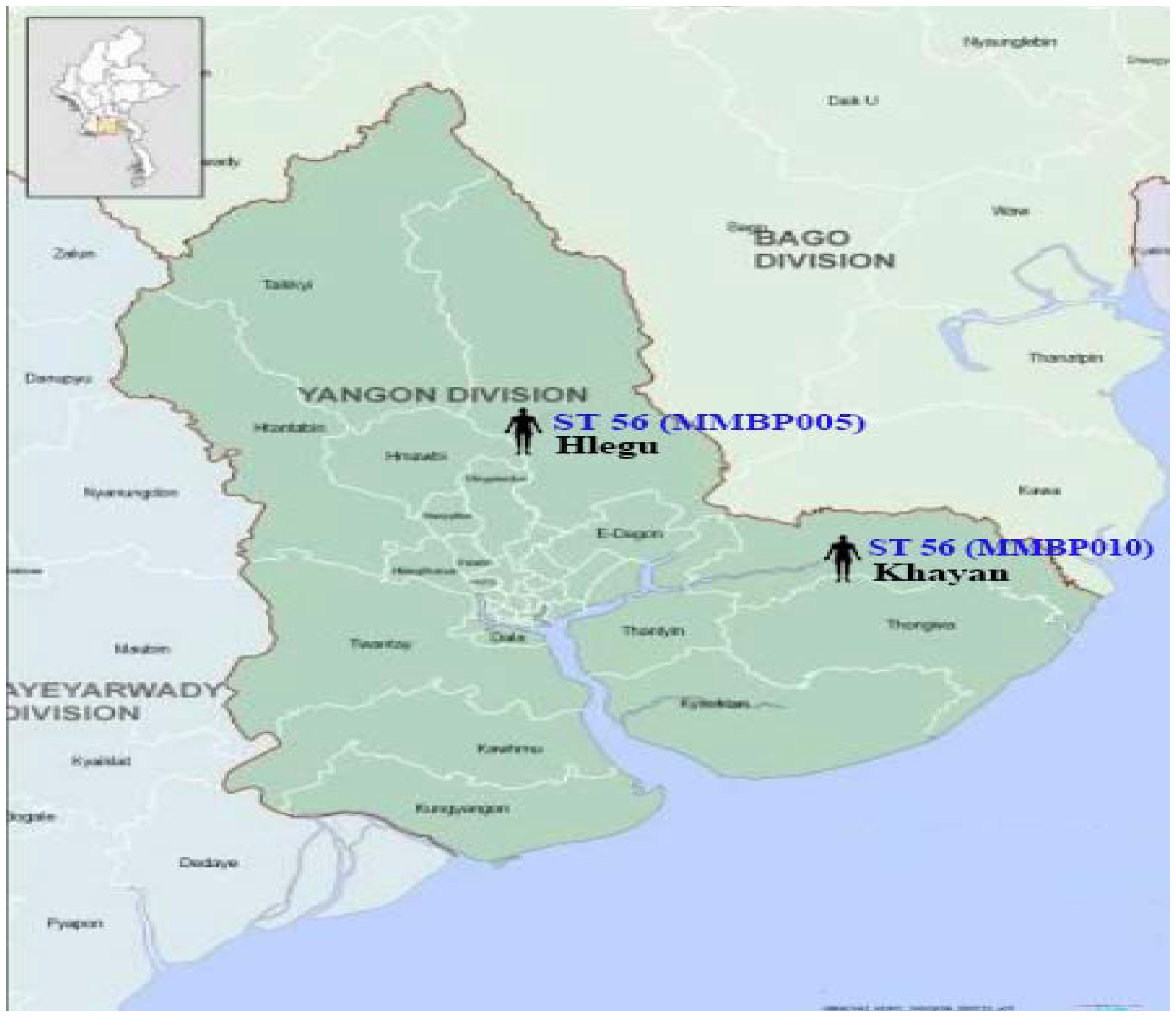
Distribution of STs 56 in Yangon division.

## Discussion

MLST is a flexible and powerful epidemiological tool to study the distribution and evolution of bacterial populations (7). ERIC PCR remains a rapid technique, easy to use, and cheap with an acceptable outcome. However, it was still problematic in its reproducibility. However, the quick assessment of *B. pseudomallei* was still essential due to its usefulness for molecular epidemiology investigations in outreach areas and low-resource countries (8). In this study, it was evaluated whether it was helpful to discriminate among STs of *B. pseudomallei*. It was likely that it could identify shared groups among the same STs. Most major band patterns of ERIC PCR exhibited approximately 80% similarity among historical STs and novel STs of the present study.

This study found two isolates (MMBP005 and MMBP010) as a single genotype in the ERIC PCR banding pattern. Surprisingly, those two isolates were isolated from different hospitals with different regions but the same province and probably infected through traveling. There was no assessment of STs from soil isolates, but an additional study should be conducted for epidemiological research in the environmental association. Antonov et al. said that ribotyping and pulsed-field gel electrophoresis are time-consuming and technically challenging for many laboratories. ERIC PCR can be used for the rapid discrimination of *B. mallei* and *B. pseudomallei* strains (9). In addition, detecting genetically diverse strains within a single geographical area highlights the complex epidemiology of B. pseudomallei and the need for continued surveillance and investigation of this pathogen in Myanmar.

For ST 90, two soil isolates were collected from the same region, but some clinical isolates were distinct and showed the same clade. Interestingly, ST 90, which was observed to be a part of a clade with one novel ST (ST 1726), exhibited approximately 80% similarity in the ERIC pattern. This finding suggests that these strains may have a common ancestor and may be related to each other. The presence of satellite variants in the MLST scheme for shared group STs (e.g., ST 1722 and ST 90, and ST 354 and ST 1725) further supports the idea of genetic diversity within these groups.

On the other hand, the novel ST 1729 was identified as an out-group in the ERIC pattern, indicating that this strain may be genetically distinct from the different strains studied. Further analysis is needed to determine the significance of this observation. A few isolates that showed a single genotype in the present study were not representative of discrimination of *B. pseudomallei*, and it pointed out for further research.

## Conclusion

This study showed that ERIC PCR represents a valuable and cost-effective alternative to MLST for molecular typing of B. pseudomallei in resource-limited settings. Its simplicity and low cost make it an attractive option for surveillance and outbreak investigations, particularly in endemic areas with limited advanced molecular methods.

## Acknowledgments

We are grateful to all the laboratory practitioners in Myanmar who collected and stored the leftover samples. We express our gratitude to Mahidol Neighboring Countries Grant’s support for this publication. We are obligated to our colleagues from the Department of Microbiology, Faculty of Medicine Siriraj Hospital, Mahidol University.

